# Oral iron exacerbates colitis and influences the intestinal microbiome

**DOI:** 10.1101/385997

**Authors:** Awad Mahalhal, Jonathan M. Williams, Sophie Johnson, Nicholas Ellaby, Carrie A. Duckworth, Michael D. Burkitt, Xuan Liu, Georgina L. Hold, Barry J. Campbell, D. Mark Pritchard, Chris S. Probert

## Abstract

Inflammatory bowel disease (IBD) is associated with anaemia and oral iron replacement to correct this can be problematic, intensifying inflammation and tissue damage. The intestinal microbiota also plays a key role in the pathogenesis of IBD, and iron supplementation likely influences gut bacterial diversity in patients with IBD. Here, we assessed the impact of dietary iron, using chow diets containing either 100, 200 or 400 ppm, fed *ad libitum* to adult female C57BL/6 mice in the presence or absence of colitis induced using dextran sulfate sodium (DSS), on (i) clinical and histological severity of acute DSS-induced colitis, and (ii) faecal microbial diversity, as assessed by sequencing the V4 region of *16S* rRNA. Increasing or decreasing dietary iron concentration from the standard 200 ppm exacerbated both clinical and histological severity of DSS-induced colitis. DSS-treated mice provided only half the standard levels of iron *ad libitum* (i.e. chow containing 100 ppm iron) lost more body weight than those receiving double the amount of standard iron (i.e. 400 ppm); p<0.01. Faecal calprotectin levels were significantly increased in the presence of colitis in those consuming 100 ppm iron at day 8 (5.94-fold) versus day-10 group (4.14-fold) (p<0.05), and for the 400 ppm day-8 group (8.17-fold) versus day-10 group (4.44-fold) (p<0.001). In the presence of colitis, dietary iron at 400 ppm resulted in a significant reduction in faecal abundance of Firmicutes and Bacteroidetes, and increase of Proteobacteria, changes which were not observed with lower dietary intake of iron at 100 ppm. Overall, altering dietary iron intake exacerbated DSS-induced colitis; increasing the iron content of the diet also led to changes in intestinal bacteria diversity and composition after colitis was induced with DSS.

## Introduction

Inflammatory bowel disease (IBD) is characterised by chronic inflammation of the gastrointestinal tract. Inflammation is associated with intestinal ulceration in both ulcerative colitis (UC) and Crohn’s disease (CD). Bleeding and malabsorption may also occur in IBD ^1,2^, and iron deficiency anaemia occurs in one-third of patients ^1, 3^. The best way to administer iron replacement to patients with IBD is a subject of debate, with both oral iron and intravenous (IV) iron supplements being considered effective ^4,5^. However, ferrous forms of oral iron replacement appear to be poorly absorbed, and the resultant free luminal iron likely results in enhanced catalytic activity and production of reactive oxygen species within the intestine ^6, 7^. High dose oral iron consumption appears to be associated with more side effects than half of the standard dose of iron ^8^, perhaps as a result of unabsorbed iron reaching the colon. Intravenous (IV) iron therapy offers effective alternative management of iron deficiency anaemia. While the route of administration is not thought to influence disease activity; oral iron supplements have been shown to disturb the microbiota, with disturbances in bacterial phylotypes and associated aberrations in faecal metabolites compared with IV treatment ^9,10^.

The gut microbiota typically comprises greater than 10^11^ microorganisms per gram of intestinal content ^11^, playing an important role in the maintenance of gut health, including protection against pathogens (colonisation resistance) and the synthesis of beneficial short-chain fatty acids (SCFA) generated through fermentation of dietary fibre ^12,13^. IBD is associated with a perturbation of gut microbiota (‘dysbiosis’), with the observed reduction in microbial diversity, including a decline in beneficial bacteria from the phyla Bacteroidetes and Firmicutes, although within these classifications a much more complicated picture exists, alongside enhancement of some potentially harmful Proteobacteria, particularly within the family *Enterobacteriaceae* ^14, 15^. The key mechanisms responsible for the development of this dysbiosis and its contribution to IBD are to date poorly defined.

Iron is an essential metal that is required by most organisms ^16^. It is a growth-limiting nutrient for many gut bacteria, which compete for unabsorbed dietary iron in the colon ^17^. Lactobacilli, considered to be beneficial intestinal barrier-maintaining bacteria, playing a significant role in the inhibition of mucosal colonisation by enteric pathogens, do not require iron ^18^. For other bacteria, acquisition of nutrient iron is an essential step for expression of key virulence factors, including Gram-negative enteric pathogens within the family *Enterobacteriaceae,* such as *Salmonella* spp., *Shigella* spp. and *Escherichia coli* pathovars ^19^. Consequently, an increase in unabsorbed dietary iron could favour the growth of opportunistic pathogens over mucosal barrier-maintaining species and alter the composition of the intestinal microbiota ^20^. In the context of IBD, excess colonic iron concentrations can occur as a result of ulceration and bleeding, as well as excess unabsorbed iron from oral iron replacement therapy ^21^.

Iron supplementation, either oral or delivered intravenously, has been shown to have an impact on the intestinal microbiota and metabolome of patients with IBD ^10^. Comparison of oral versus intravenous routes of iron supplementation demonstrated no significant effects on the human bacterial diversity, although specific species changes were noted. Four Operational Taxonomic Units (OTUs) were observed to be less abundant after oral iron therapy, including *Faecalibacterium prausnitzii,* low abundance of which, has been linked to relapse in Crohn’s disease ^10^. An OTU of the *Bifidobacterium* genus was noted to be increased with oral iron therapy but, the effect of prebiotics and probiotics was a confounder in 4/6 patients studied. Overall, this study suggested that oral iron therapy might have an adverse effect on the microbiome; however, the contemporaneous effect of IBD was not studied.

Murine models of IBD offer the opportunity to investigate the gut microbiota and microbial diversity changes that occur during IBD pathogenesis ^22^, ^23^. We hypothesised that changing the amount of dietary iron would influence IBD development in a murine model of colitis. Hence, in this study, we modified the standard chow diet of C57BL/6 mice (at 200 parts per million-ppm iron) to half of the standard levels (100 ppm), or to double that of standard (400 ppm) of iron and subsequently induced colitis using dextran sulfate sodium (DSS). Clinicopathological outcomes were analyzed in parallel with the characterisation of the composition of the gut microbial community by sequencing the *16S* prokaryotic ribosomal subunit.

## Materials and Methods

### Animals

Female C57BL/6 mice (n=130), aged 8-9 weeks old, were purchased from Charles River Laboratories (Margate, UK). Mice were fed a standard rodent chow pellet diet, during an initial acclimatisation period of at least one week, with access to water *ad libitum.* All mice were individually-caged in a specific pathogen-free animal facility with controlled temperature, humidity and a pre-set dark-light cycle (12 h: 12 h). Eight groups were studied initially; two control groups and six DSS-treated groups were maintained either for 8 days or up to 10 days. Three additional control groups of mice (to compare the effects of diets alone) received drinking water without DSS, but with varying amounts of dietary iron for a total of 10 days, under conditions as described above (see Table 1). For each set of experiments, mice were matched for age and body weight. The work described was conducted in accordance with UK Home Office regulations under the Animals (Scientific Procedures) Act 1986 (ASPA). The University of Liverpool Ethical Review Body also approved protocols. Study animals were observed for signs of illness and/or welfare impairment and were euthanised by cervical dislocation.

**Table 1:**
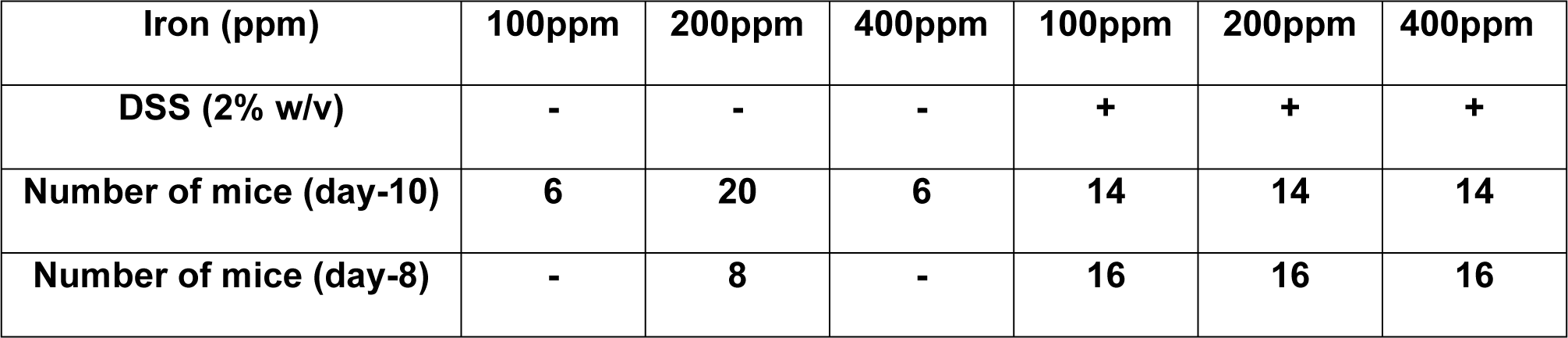
Experimental animal groups’ classification

### Diets

The standard 10 mm compression pellet chow diet utilised contained 200 ppm iron (Rat and Mouse Breeder and Grower Pelleted CRM (P) - Special Diets Services, Witham, Essex, UK). Two modifications of this diet were used. The first diet was formulated to contain half the amount of iron found in standard chow, ie. 100 ppm, a dietary level selected to reduce luminal bacterial exposure to iron without being harmful to the mice. The second formulation contained double the amount of iron found in standard chow, i.e. 400 ppm, a diet to increase bacterial exposure to iron without being overtly toxic to mice.

### Induction of acute colitis using dextran sulfate sodium (DSS)

Mice were given 2% w/v dextran sodium sulfate (M.W. 36,000 - 50,000Da; Catalogue number: 160110; Lot number: 6683K; MP Biomedicals, UK) in their drinking water for 5 days to induce colitis (~150 mL/mouse over 5 days), followed by another 5 days of DSS-free drinking water. Mice were euthanised on day-8 or day-10.

### Histopathological scoring of colonic inflammation

The distal colon was removed, fixed in 4μ neutral buffered formalin, dehydrated, paraffin wax-embedded and then 4 μm sections were cut by microtomy. The sections were stained with hematoxylin and eosin (H&E), and inflammation scored, using the system described by Bauer *et al.* ^24^.

### Measurement of faecal calprotectin as a marker of the degree of intestinal inflammation

Faecal pellets were collected from each cage (1 mice per cage), in all groups, on day 1,8 and 10. Faecal calprotectin levels were measured using an S100A8/S100A9 ELISA kit (Immundiagnostik AG, Bensheim; Germany) as per the manufacturer instructions.

### Assessment of faecal iron

The faecal iron (Fe^2+^ and Fe^3+^) concentration was measured using an iron immunoassay kit [MAK025, Sigma-Aldrich]. This was performed using faecal pellets taken at the same time as those for the faecal calprotectin ELISA.

### High-throughput sequence analysis of bacterial communities from faecal samples

Faeces (2 g) was sampled from each animal and bacterial DNA extracted using the Stratec PSP^®^ Spin Stool DNA Plus Kit (STRATEC Molecular GmbH, Berlin; Germany) following the manufacturer recommended protocol. Isolated DNA was sent to the Centre for Genomic Research at the University of Liverpool to generate the *16S* Metagenomic Sequencing Library. Primers described by Caporaso *et al.^25^* were used to amplify the V4 region of *16S* rRNA; F: 5’ACACTCTTTCCCTACACGACGCTCTTCCGATCTNNNNNGTGCCAGCMGCCGCGGT AA3’ and R: 5’GTGACTGGAGTTCAGACGTGTGCTCTTCCGATCTGGACTACHVGGGT WTCTAAT3’.

Approximately 5 μL of extracted DNA was used for first round PCR with conditions of 20 sec at 95°C, 15 secs at 65°C, 30 sec at 70°C for 10 cycles then a 5 min final extension at 72°C. Amplicons were purified with Axygen SPRI Beads before a second-round PCR was performed to incorporate Illumina sequencing adapter sequences containing indexes (i5 and i7) for sample identification as described in ^25^. Fifteen cycles of DNA amplification by PCR were performed using the same conditions as above, i.e., 25 cycles overall. Again, samples were purified using Axygen SPRI Beads before being quantified using Qubit and assessed using the Fragment Analyser. Successfully generated amplicon libraries were used for sequencing.

The final libraries were pooled in equimolar amounts using the Qubit and Fragment Analyser, data and size-selected on the Pippin Prep using a size range of 350-550 base pairs (bp). The quantity and quality of each pool were assessed by Bioanalyzer, and subsequently by qPCR using the Illumina Library Quantification Kit from Kapa on a Roche Light Cycler LC480II system according to the manufacturer instructions. The pool of libraries was sequenced on one lane of the MiSeq at 2 × 250 bp paired-end sequencing ^26^. To help balance the complexity of the amplicon library, 15% PhiX was spiked as described by Altschul *et al.* ^27^.

### Bioinformatics analysis

Initial processing and quality assessment of the sequence data was performed using an in-house pipeline. Base calling and de-multiplexing of indexed reads were conducted by CASAVA version 1.8.2 (Illumina) as described by Schubert *et al.* ^28^. The raw fastq files were trimmed to remove Illumina adapter sequences with any reads matching the adapter sequence over at least 3 bp being trimmed off. The reads were further trimmed to remove low-quality bases (reads <10 bp were removed). Read pairs were aligned to produce a single sequence for each read pair that would entirely span the amplicon. Sequences with lengths outside of the expected range (which are likely to represent errors) were also excluded. Sequences passing the above filters for each sample were pooled into a single file. A metadata file was created to describe each sample. These two files were used for metagenomics analysis using Qiime, version 1.8.0 as described by Caporaso *et al.* ^29^. Similar sequences were clustered into groups, to define OTUs of 97% similarity. OTU-picking was performed using USEARCH7 as described by Edgar *et al.* ^30^ to cluster sequences, remove chimeras, and define OTU abundance. The Greengenes database of ribosomal RNA sequences, version 12.8 as described by McDonald *et al.* ^31^, was used as a reference for reference-based chimera detection. To reduce the effect of sample size and to estimate species richness within each sample (alpha diversity), OTU tables were repeatedly sub-sampled (rarefied). For each rarefied OTU table, three measures of alpha diversity were estimated: Chao1, the observed number of species, and the phylogenetic distance. To allow inter-sample comparisons (beta-diversity), all datasets were sub-sampled (rarefied). Rarefied OTU tables were used to calculate weighted and unweighted pair-wise UniFrac matrices. UniFrac matrices were then used to generate UPGMA (Unweighted Pair-Group Method with Arithmetic mean) trees and 2D principal component analysis (PCA) plots.

### Statistics

Normally distributed physiological and biochemical data (as determined by Shapiro-Wilks test) were assessed by one-way analysis of variance followed by multiple pairwise comparisons of treatment means using Dunnett’s test. Non-normally distributed data were assessed by Kruskal-Wallis non-parametric-test followed by multiple pairwise comparisons (Conover-Inman) test (Stats Direct version 3.0.171; Altrincham, UK).

For the bioinformatic analysis of microbiota data, Welch’s t-test was used with the false discovery rate (FDR) Storey’s multiple correction tests. The q-value is the adjusted p-value based on FDR calculation, where statistical significance was declared at p<0.05.

## Results

### Reduced dietary iron intake is associated with increased weight loss and more severe colitis following the induction of DSS colitis

All mice treated with 2% w/v DSS lost body weight from day-5, with maximal weight loss occurring on day-8. Mice ingesting a diet containing 100 ppm iron lost significantly more weight (13 ± 1.53%) than seen with the other DSS treatment groups (i.e. 8.3 ± 1.09% for mice ingesting 200 ppm iron, and 8.6 ± 1.33% for mice on 400 ppm iron); see Fig 1. Control mice, ingesting dietary iron at 100 ppm, 200 ppm and 400 ppm, receiving no DSS treatment, showed expected steady increases in body weight over the 10-day study period. No evidence of colitis was observed in all untreated (controls) mice. In contrast, all mice treated with 2% w/v DSS developed bloody diarrhoea within the last 5 days of the 10-day study. Histopathological examination established the presence of DSS-induced colitis, which was localised mainly to the distal part of the colon. Histological features of colitis observed included areas of mucosal loss, inflammatory cell infiltration and oedema (Fig 2). Histological colonic inflammation severity scores were significantly greater in mice consuming 100 ppm dietary iron and treated with 2% w/v DSS, compared with those mice receiving 2% w/v DSS and ingesting a diet containing 200 ppm or 400 ppm iron, at both day-8 and day-10 (Fig 3).

**Fig 1:**
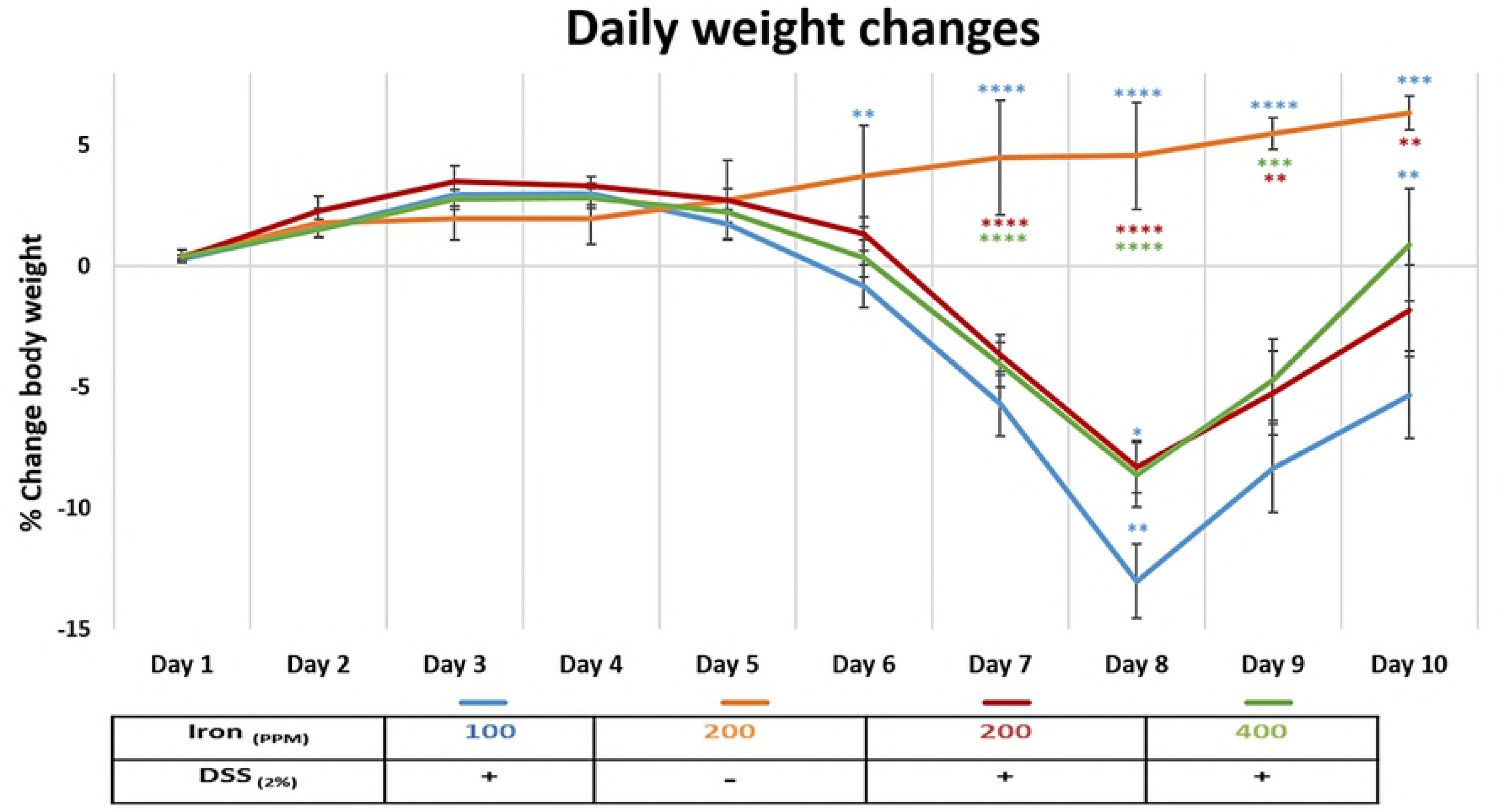
This is the Fig 1 Title: Daily weight changes. This is the Fig 1 legend: Percentage weight change in mice consuming diets containing iron [100 ppm (blue), 200 ppm (red) and 400 ppm (green)] during dextran sulfate sodium (DSS)-induced colitis, and mice consuming a diet containing 200 ppm iron without DSS treatment (orange) during the 10-day study period. Data are presented as a mean ± standard error of the mean (SEM). Statistical differences were assessed by Kruskal-Wallis test followed by multiple comparisons (Conover-Inman) tests (* p<0.05, ** p<0.01, *** p<0.001, **** p<0.0001). (n=30 female mice per DSS-treated; n=22 mice per untreated group).

**Fig 2:**
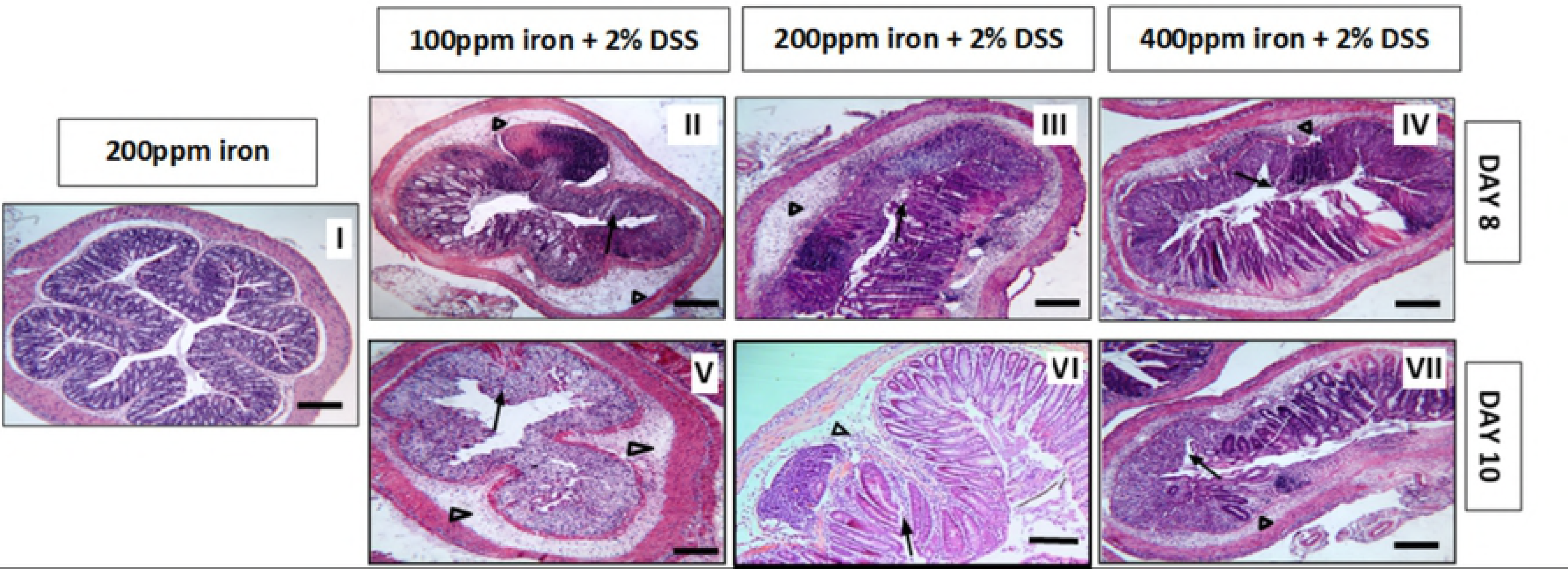
This is the Fig 2 Title: H & E histology. This is the Fig 2 legend: Representative Haematoxylin- and eosin-stained sections of the distal colon from untreated and 2% w/v DSS-treated mice. Mice received either water (control, I) or 2% w/v DSS for 5 days followed by another 3 days on plain drinking water (II, III and IV) or 5 days on plain drinking water (V, VI, and VII). Arrowheads highlight submucosal oedema; arrows highlight almost complete loss of colonic epithelium.

**Fig 3:**
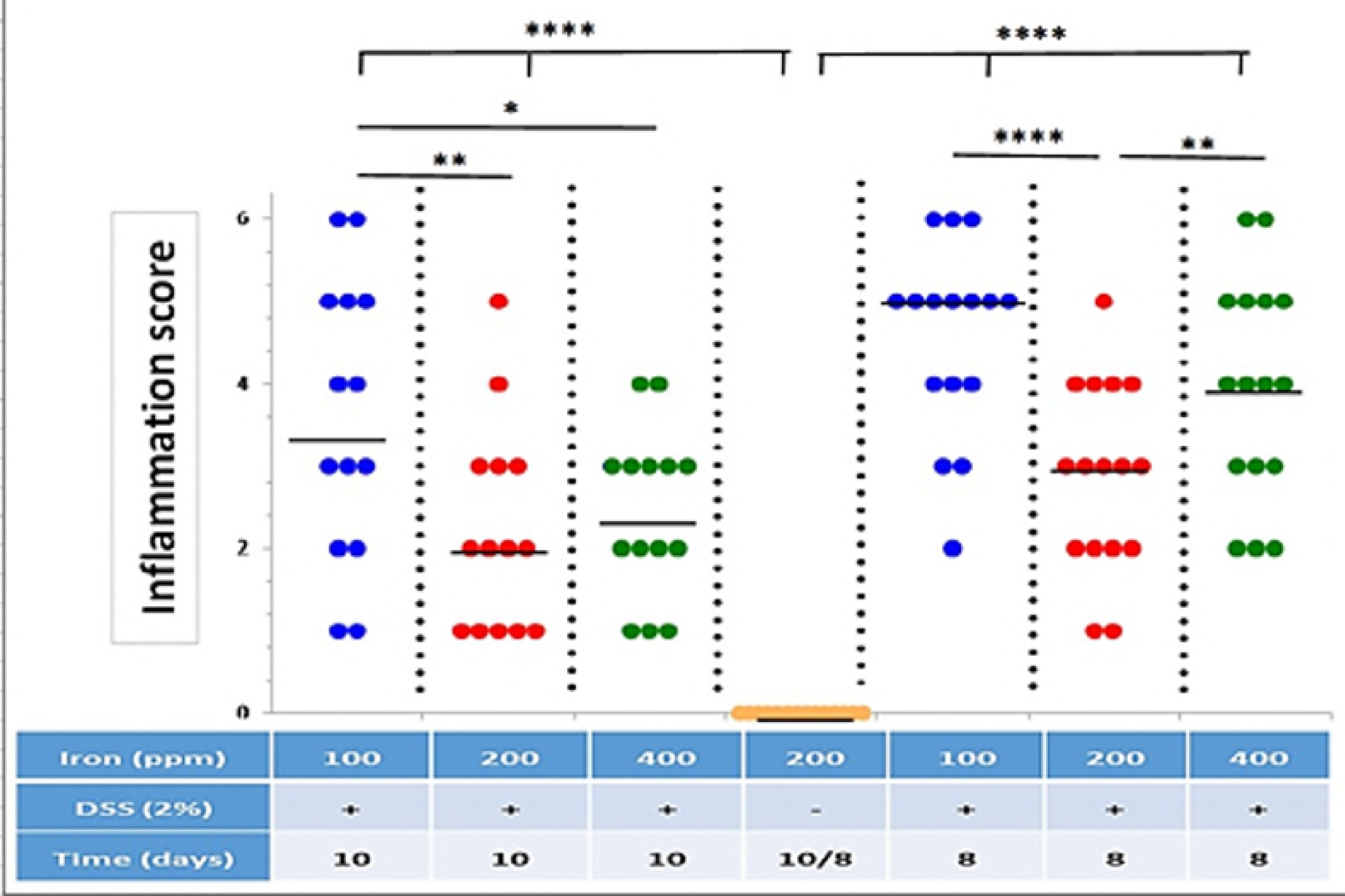
This is the Fig 3 Title: Inflammation score. This is the Fig 3 legend: Inflammation (colitis) scores for all groups of DSS-treated mice (n=16 [8-days] and n=14 [10-days] mice per group) and untreated control mice (n=24) on diets containing different levels of iron (100, 200 and 400 ppm). Horizontal lines represent medians. Significant differences were assessed using one-way ANOVA followed by multiple comparisons against untreated control by Dunnett’s test; * p<0.05, ** p<0.01, **** p<0.0001.

### Faecal calprotectin concentration in DSS-treated mice during the 10-day course

Faecal calprotectin concentrations were measured in faecal pellets collected from the cage of each mouse in all groups at day-1, day-8, and day-10. Analysis of day-1 samples showed similar levels (10.3 ‘mean’) in all groups indicating no treatment effects were yet apparent. Faecal calprotectin concentration data were normalised to the values found in control samples with higher levels seen in the mice on modified (half and double of the standard content) iron diet compared to those fed the standard 200 ppm iron diet (Fig 4). This finding was seen at both day-8 and day-10 although the most striking levels were recorded at day-8 (60 ± 1.11%, 40 ± 1.12% and 80 ± 1.08% increase for the half of the standard, standard and the double of the standard iron diets, respectively). The maximal faecal calprotectin levels were seen at day-8 and correlated with the highest histological scores in DSS-treated mice that received 200 ppm and 400 ppm iron containing diets compared to their corresponding day-10 non-DSS controls (p<0.001).

**Fig 4:**
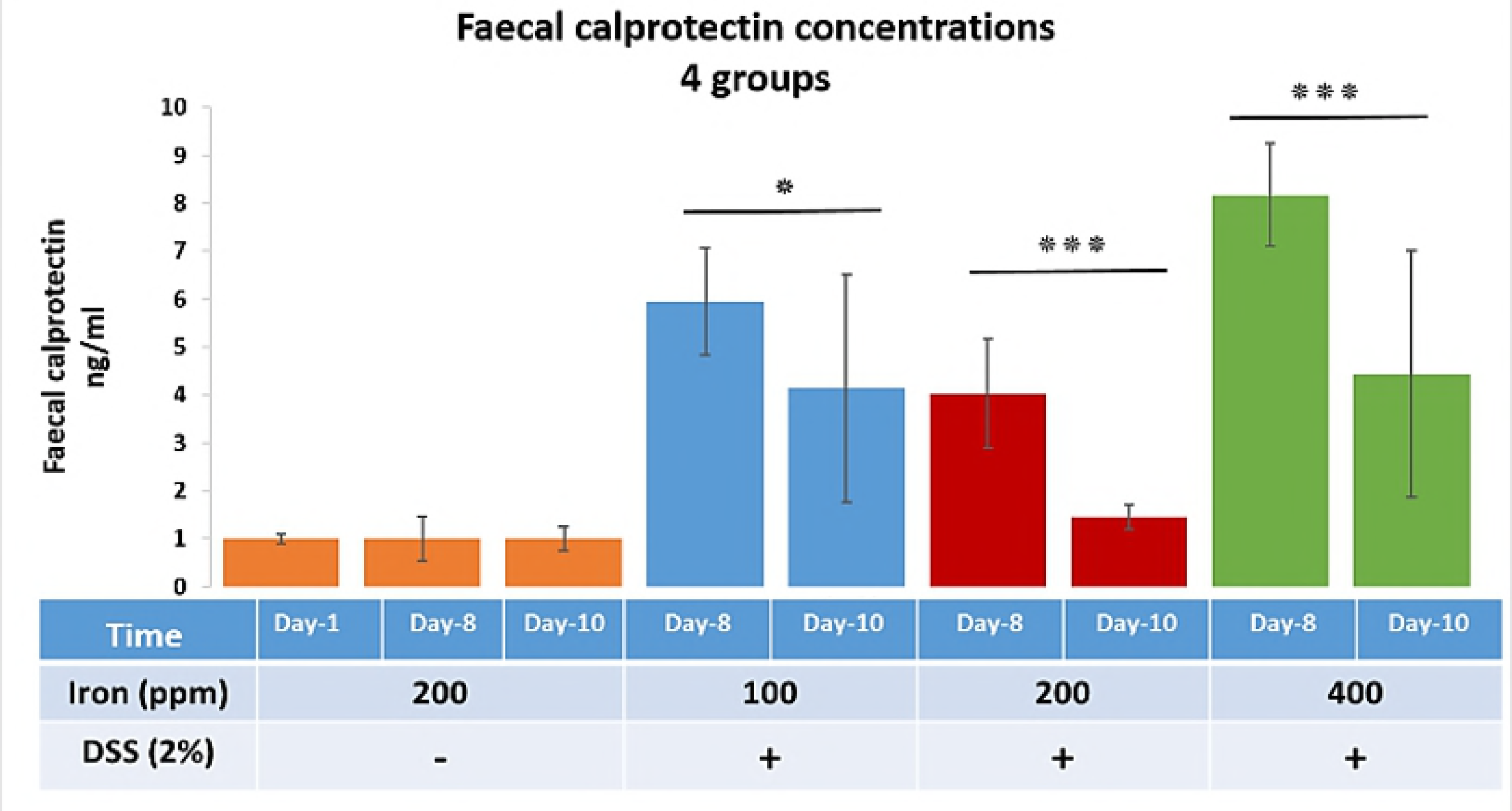
This is the Fig 4 Title: Faecal calprotectin concentrations. This is the Fig 4 legend: Faecal calprotectin concentrations at three different time points (day-1, 8 and 10) for four groups, three DSS-treated groups (consuming diets containing 100, 200 and 400 ppm iron) and one untreated control group (consuming a standard 200-ppm iron containing chow diet). Data are presented as a mean ± SEM. Significant differences were identified using the Kruskal-Wallis test followed by multiple comparisons (Conover-Inman) test; * p<0.05, *** p<0.001. (30 samples for all DSS-treated groups and 22 samples for untreated mice at each time point).

### Faecal iron concentrations in DSS-treated mice

Faecal iron concentrations were measured to investigate the net impact of dietary iron and bleeding resulting from inflammation. Faecal pellets, from each mouse, were assessed for faecal iron concentration (ferric and ferrous) from any (dietary and bleeding) source at the different time points (day-1, 8 and 10). Data points from experimental groups were normalised to the values found in the control samples (Fig 5). Faecal iron concentrations were increased, at day-10, for all mice with DSS-induced colitis, compared to control mice, with the greatest level of change being observed in mice receiving half of standard chow dietary iron levels, i.e. 100 ppm (Fig 5). DSS-treated mice receiving the standard levels of iron (200-ppm diet) had significance (P<0.05) faecal iron concentrations at day-8 vs day-8 within the control group of mice. Observed differences in faecal iron concentrations between mice on half of standard chow dietary iron levels (100 ppm) and double the standard iron diet levels (400 ppm) were not statistically significant. This suggests that colitis and bleeding likely had more pronounced effects on the faecal iron concentration than the amount of iron consumed in the diet alone.

**Fig 5:**
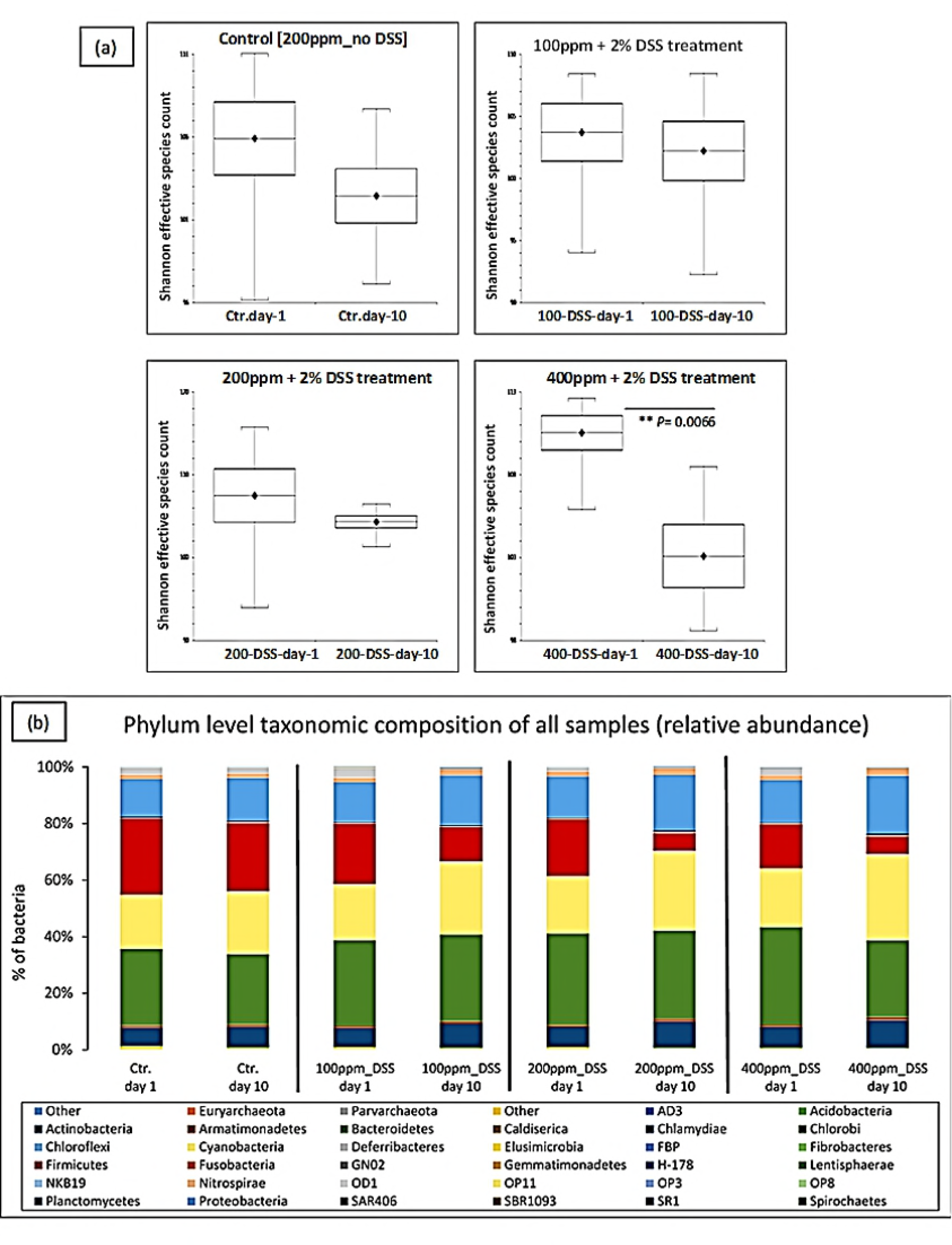
This is the Fig 5 Title: Faecal iron concentrations. This is the Fig 5 legend: Faecal iron concentration at three different time points (day-1, 8 and 10) for four groups, three DSS-treated groups (consuming diets containing 100, 200 and 400 ppm iron) and one untreated control group (consuming a standard 200-ppm iron containing chow diet). Data are presented as a mean ± SEM. Significant differences were identified using the Kruskal-Wallis test followed by multiple comparisons (Conover-Inman) test; * p<0.05.

### Effect of iron on the microbiota composition in the colon after DSS-induced colitis

To determine the effect of DSS and oral iron on the gut microbiota, fresh faecal samples were compared at baseline and the end of each experiment (after 10 days from the start). After sequence processing and filtering, a total of 11,811,301 chimera-checked *16S* rRNA sequences (166,356 ± 59,353 per sample) spanning a total of 204,331 OTUs were obtained.

Analysis of alpha-diversity (statistical significance of Shannon) indicated that there was a significant reduction in species richness in faecal samples taken from the 400 ppm iron fed, the DSS-treated group between day-1 and day-10 (P<0.0066; Shannon diversity index) (Fig 6-a). To assess whether the differences in species richness were attributable to alterations in the relative abundance of specific bacterial groups, we compared the proportions of various taxonomic groups at the phylum level. Bacteroidetes was the most abundant phyla present, followed by Firmicutes, Cyanobacteria and Proteobacteria (Table 2). Phyla changes were seen in all DSS-treated groups when day-10 samples were compared to day-1. However, these changes were only observed to be statistically significant for the mice consuming 400 ppm iron, with increases observed in the numbers of Proteobacteria (increased 1.40 ± 0.1-fold) and Actinobacteria (1.30 ± 0.1-fold increase) and concomitant reductions in Firmicutes (0.6 ± 0.1-fold) and Bacteroidetes (0.8 ± 0.04-fold); these changes have been explicitly accredited to Proteobacteria, Actinobacteria, Firmicutes and Bacteroidetes, which occurred in the presence of inflamed versus non-inflamed tissues even within the same group. Therefore, the double of the standard iron diet group had the highest relative abundance among inflamed (colitis) groups on the subject of reduction or increase changes (day-1 vs day-10) (Table 2 and Fig 6-b).

**Table 2:** **Comparison between all groups regarding proportions of bacteria at the phylum level at day-1 vs day-10**

| Taxonomy | Control Day-1 | Control Day-10 | 100ppm DSS Day-1 | 100ppm DSS Day-10 | 200ppm DSS Day-1 | 200ppm DSS Day-10 | 400ppm DSS Day-1 | 400ppm DSS Day-10 |
| --- | --- | --- | --- | --- | --- | --- | --- | --- |
| Actinobacteria | 6.63% | 7.25% | 6.77% | 8.48% | 7% | 9.28% | 7.35% | 9.82% |
| Bacteroidetes | 27.03% | 24.98% | 30.38% | 30.73% | 32% | 31.33% | 34.62% | 27.27% |
| Cyanobacteria | 19.13% | 22.02% | 19.80% | 25.67% | 20% | 27.87% | 20.70% | 30.23% |
| Firmicutes | 26.98% | 24.15% | 21.33% | 6.20% | 20% | 10.2% | 15.57% | 6.22% |
| Proteobacteria | 13.22% | 15.17% | 13.85% | 17.55% | 15% | 19.65% | 14.82% | 20.58% |
| TM7 | 1.63% | 1.63% | 1.65% | 2.02% | 2% | 2.27% | 1.80% | 2.48% |
| Tenericutes | 2.42% | 1.90% | 3.55% | 0.45% | 1% | 0.15% | 2.53% | 0.07% |

**Fig 6:**
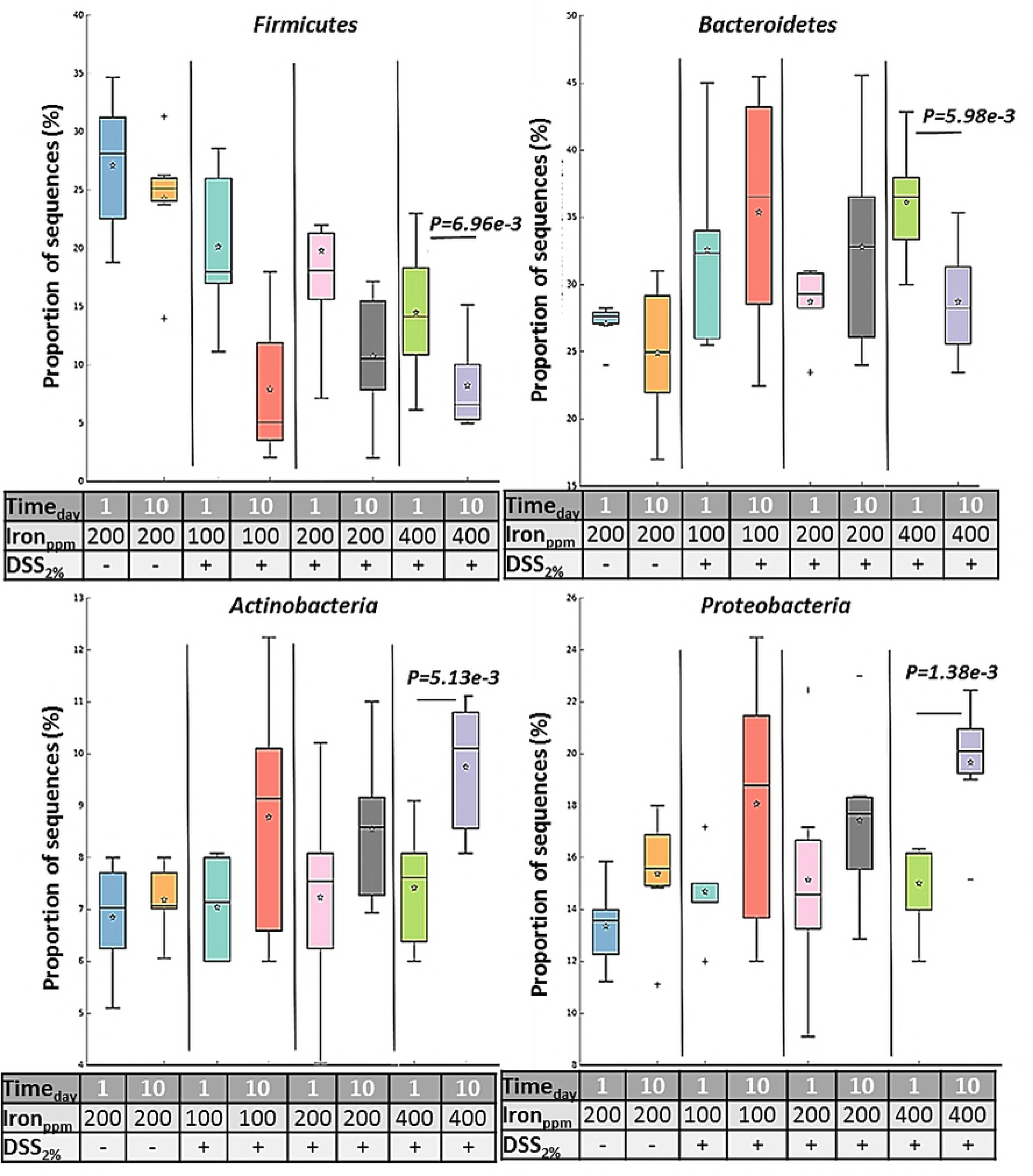
This is the Fig 6 Title: Relative abundance of bacteria. This is the Fig 6 legend: Effect of iron on the microbiota composition in the colon after DSS-induced colitis. (a) Shannon effective diversity boxplots display decreased numbers of dominant molecular species in all groups, day-1 versus day-10 of the study. (b) The Phylum-level taxonomic composition of all samples (average relative abundance). Ctr. = untreated controls on a standard chow diet containing 200 ppm iron; DSS = 2% w/v dextran sulfate sodium (DSS) treated mice on diets containing low iron (100 ppm), standard iron (200 ppm) and high iron (400 ppm) levels.

We searched for differences between day-1 and day-10 samples by considering delta-values calculated as differences in sequence abundances (before and after treatment). No, statistically significant changes were observed in mice receiving diets containing half of the standard iron levels where DSS was administered, despite showing similar trends to those mice on double the standard diet iron levels (Fig 7).

**Fig 7:**
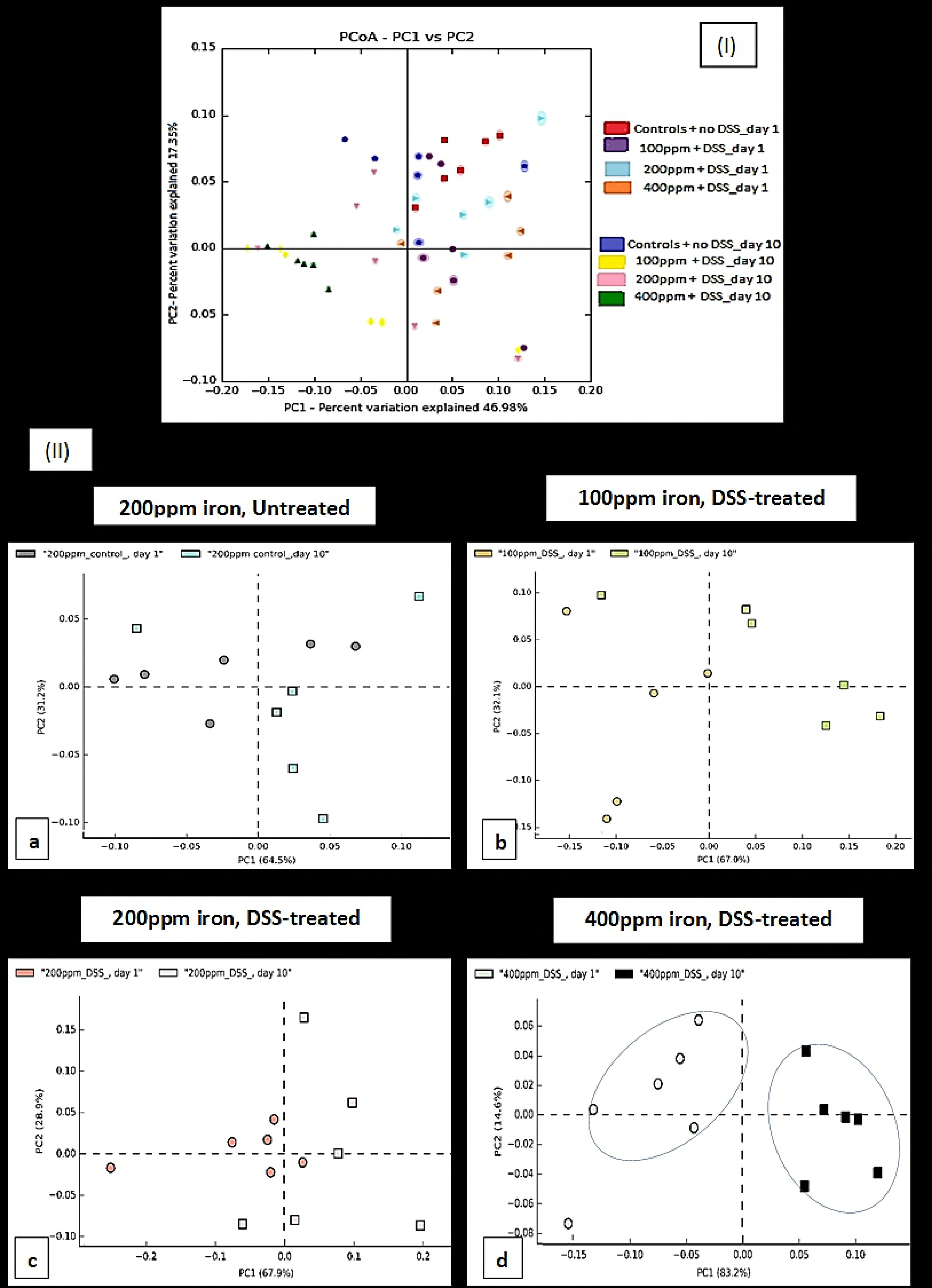
This is the Fig 7 Title: Proportions of sequences. This is the Fig 7 legend: Box plot is showing the distribution in the proportion of four key phyla (Firmicutes, Bacteroidetes, Actinobacteria and Proteobacteria) assigned to samples from all groups at day-1 and day-10. Boxes indicate the interquartile ranges (75^th^ to 25^th^ IQR) of the data. The median values are shown as lines within the box, and the mean values are indicated by stars. Whiskers extend to the most extreme value within 1.5*IQR. Outliers are shown as crosses. Statistical differences were assessed by Welch’s t-test followed by Storey’s FDR multiple test correction.

Principal Component Analysis (PCA) was used to identify linear combinations of gut microbial taxa that were associated with specific diets. There was a clear separation of samples from the mice consuming a chow diet containing 400 ppm before (day-1) and after (day-10) DSS-treatment which was not seen in the other treatment groups (Fig 8). This suggests that DSS-induced colitis, in the presence of double the standard level of dietary iron intake, affected the bacterial community significantly more than that observed in all other diet groups (P<0.0066; Shannon diversity index) (Fig 6-a).

**Fig 8:**
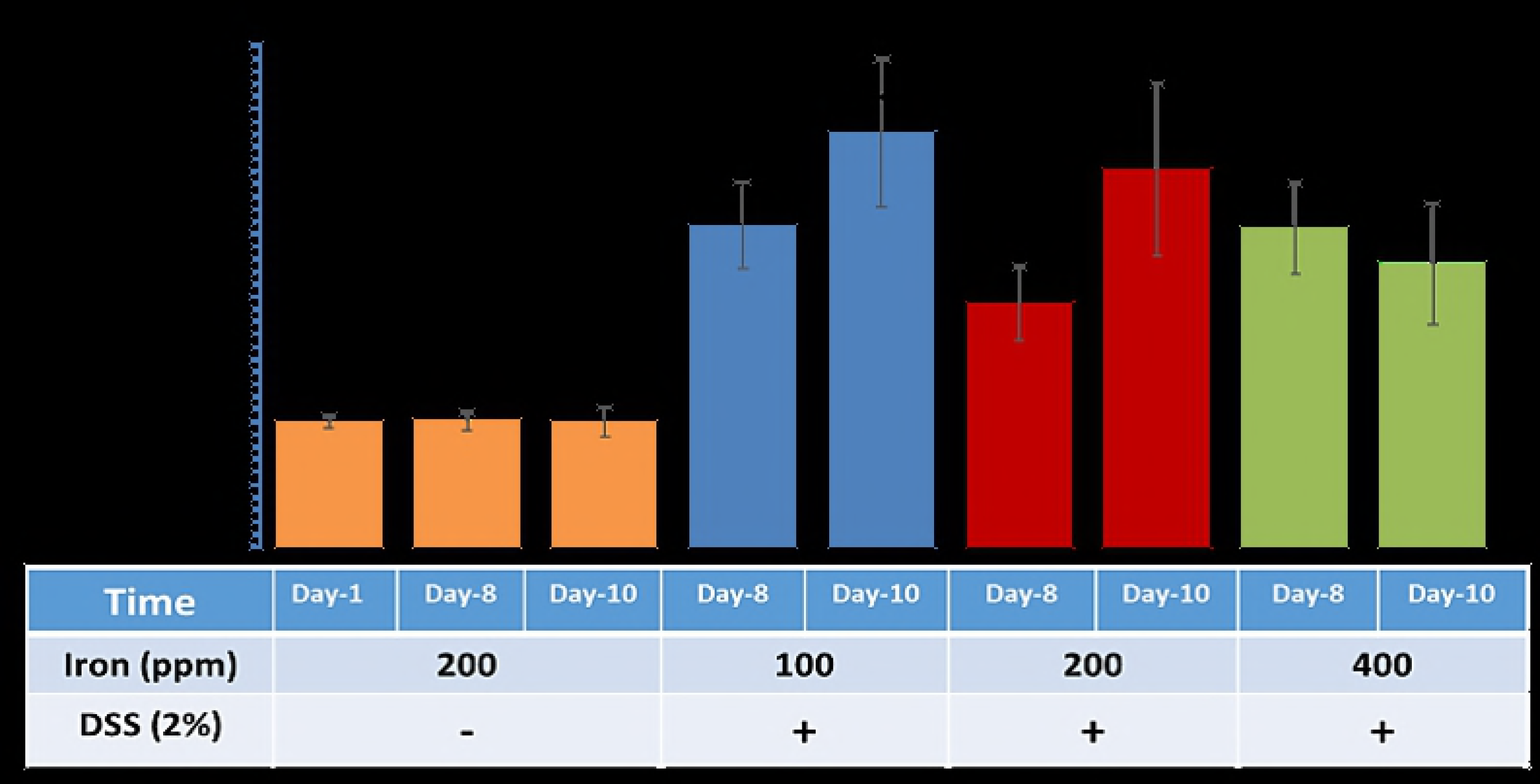
This is the Fig 8 Title: Principal Coordinate/Component Analysis. This is the Fig 8 legend: Analysis of faecal microbiota shifts assessed by Principal Coordinate/Component Analysis (PCA-PCoA) plots of the unweighted UniFrac distances of pre-and post-DSS-intervention stool samples (I) PCoA ; all groups (II) PCA; DSS-treated mice on diets containing low iron (100 ppm), standard iron (200 ppm) and high iron (400 ppm) (b, c, and d respectively) and untreated control mice on a diet containing standard 200 ppm iron (a) at phylum-level, phylogenetic classification of *16S* rRNA gene sequences. Symbols represent data from individual mice, colour-coded by the indicated metadata. Statistical differences were assessed by Welch’s t-test followed by Storey’s FDR multiple test correction.

## Discussion

In this study, we used a murine model of IBD in which 2% w/v DSS was administered to mice and investigated the impact of changing dietary iron intake on the degree of inflammation and the bacterial components of the intestinal microbiome.

Alteration of iron content from standard chow diet levels (200 ppm) significantly influenced the severity of colitis induced by DSS in mice. Clinically, for mice treated with DSS, those fed half the standard iron levels developed more severe colitis (compared to those consuming chow diets with iron levels at 200 ppm, or at higher levels of 400 ppm. DSS-treated mice that received 100 ppm dietary iron significantly also lost more body weight than observed in the other treatment groups. However, at molecular level increasing dietary iron 2-fold above standard levels, to 400 ppm, led to worse inflammation and greater faecal calprotectin concentrations at day-8, than was found in mice consuming a 100 ppm iron diet. Our observation agrees with the findings of an earlier study performed by Carrier and colleagues ^32^ in DSS-treated rats which emphasised the role of nutrient iron in modulating inflammation. Specifically, the severity of colitis appeared to positively associate with the amount of iron consumed. However, they did not investigate the effects of consumption of lower than normal amounts of iron in their work ^32^. A study by Erichsen *et al.* ^33^ reported that the addition of low-dose oral ferrous fumarate (0.60 mg Fe/kg/d) to Wistar rats to levels present in standard chow 130 mg/kg (ferrous carbonate, 40 mg/kg; the remainder representing organic iron), also increased the severity of DSS-induced colitis. In the same study, oral supplementation with higher doses of ferrous fumarate caused a further increase in histological intestinal inflammation ^33^. Our study shows that a diet depleted in iron (100 ppm) can also exacerbate colitis severity. The mice that consumed a diet containing 100 ppm iron, and treated with 2% w/v DSS, showed greater increased intestinal inflammation than mice ingesting a standard chow diet containing 200ppm iron, and treatment with 2% w/v DSS.

It has previously been suggested that iron formulations can be beneficial (ferrous bisglycinate) or highly damaging (ferric ethylenediaminetetraacetic acid (FEDTA)) during DSS-induced colitis experiments ^9^. Iron supplementation at different doses also induced shifts in the gut microbial community and inferred metabolic pathways ^9^. Our findings indicate that any significant alteration in standard dietary iron (above or below the standard chow levels of 200 ppm) may have a negative impact on the severity of DSS-induced colitis in mice.

For humans, faecal calprotectin measurement is commonly used as an assessment tool for disease activity in IBD ^34, 35^. We, therefore, used this additional approach and measured murine faecal calprotectin levels to examine whether dietary iron levels affected inflammation. The degree of colonic inflammation was found to be significantly higher for DSS-treated mice receiving 400 ppm iron in their chow as assessed by faecal calprotectin concentration. The histopathological changes observed were consistent with the faecal calprotectin levels measured, which were higher at day-8 than at day-10, particularly in the high- and low-iron fed, DSS-treated groups. A previous study in African infants by Jaeggi and colleagues ^36^ also noted that oral iron supplementation was associated with increased concentrations of faecal calprotectin and with an increased rate of diarrhoea ^36^. In contrast, a mouse study by Kortman *et al.^37^* showed that faecal calprotectin concentrations were not influenced by dietary iron intervention alone, but only following an enteric infection *(Citrobacter rodentium),* with faecal calprotectin concentrations being significantly lower in mice consuming an iron-deficient diet. Kortman *et al.* ^37^ also found that Gram-positive *Enterorhabdus* appeared only after enteric infection and its relative abundance, and faecal calprotectin concentrations observed, were highest in a standard (45 mg/kg) dietary iron group ^37^.

In the present study, all DSS-treated mice showed an increase in faecal calprotectin levels at day-8; this was most prominent in the mice consuming 400 ppm dietary iron. However, all DSS-treated groups showed greater levels of calprotectin in their stool at day-8 vs day-10.

This further supports the view that altering the standard levels of dietary iron may exacerbate the severity of murine DSS-induced colitis.

One key source of iron accessible to the intestinal microbiota is unabsorbed, excess dietary iron and any significant changes in luminal iron concentrations may have a potential impact on structure, function and diversity of the intestinal microbiome ^36, 38^. Iron replacement therapy is a common treatment in patients with anaemia and IBD, such as in Crohn’s disease, although such supplements may also influence intestinal inflammation as well as intestinal microbial community structure and function ^32, 39^.

Measuring faecal iron concentrations would help to assess the severity of bleeding during colitis. However, it is difficult to distinguish between the iron that comes from the diet and that which has been released from red blood cells because of luminal bleeding during colitis. Following a collection of faecal pellets at different time points from each mouse and calculating the iron content (ferric and ferrous) (dietary and bleeding source) and comparing results observed between groups at day-10, the absolute amount of faecal iron appeared to be different for DSS-treated groups (3.3-fold increase in half of the standard iron group, and 2.3-fold increase in the double the standard iron group compared with the control group at day-10). There was a significant increase in faecal iron at day-8, in standard iron diet group, but not in the other groups. As there was as if an to increased (no significance) in faecal iron in mice fed 100 ppm iron compared with those mice fed the standard chow diet level of 200 ppm, this suggests that luminal bleeding may be a contributing factor to faecal iron quantitation in the DSS-induced colitis model.

Overall in this study, changes (increase or decrease) in the iron content of the diet from standard chow levels (200 ppm) appeared to significantly enhance colonic inflammation in a DSS-induced mouse model of IBD. There appeared to be synergy between dietary iron levels and DSS treatment of colonic inflammation and faecal calprotectin levels. Faecal iron concentrations are known to be increased by inflammation, as well as oral iron intake ^36^. This may explain the paradox in the half standard dietary iron fed group where luminal bleeding during colitis caused an increase in the faecal iron concentration despite lower levels of iron being consumed in the diet.

Changes in the microbiota are thought to be a major contributory factor in many human diseases, including IBD ^40 41^. The most distinct phylum level alterations in IBD are a reduction in the abundance of Bacteroidetes and Firmicutes and increased proportions of Proteobacteria, in particular, increased numbers of bacteria from the family *Enterobacteriaceae* ^14, 40, 42, 43^. Murine models of IBD provide a means to investigate bacteria in IBD ^22^, and dysbiosis of the intestinal microbiota has been shown to induce murine colitis ^23, 44^. Here, we analysed inter- and intra-group differences and similarities between the intestinal microbiota composition of 24 laboratory C57BL/6 mice (6 mice/group). Qualitative and quantitative-based analysis of the faecal gut microbiota at two different time points (day-1 and 10) for DSS-treated groups (100, 200 and 400 ppm dietary iron) and untreated mice (controls) was undertaken. Principal component analysis indicated an overlap of all microbial profiles, except for the double standard dietary iron (400 ppm) fed DSS-treated mice. Based on the PCA, ingestion of double the standard level of dietary iron was found to be the most important factor responsible for clustering.

Some studies have shown that subsets of CD and UC intestinal tissue and faecal samples have an abnormal gut microbiota, characterised by depletion of commensal bacteria, in particular members of the phyla Firmicutes and Bacteroidetes, and an increase in Proteobacteria ^14, 43^. Doubling the standard level of iron in the chow diet (i.e. to 400ppm) here led to significant alterations in microbiota composition in 2% w/v DSS-treated mice, with our study showing a similar pattern of change to those observed in human IBD, including increases in Proteobacteria and concomitant decreases in Firmicutes and Bacteroidetes.

Similar trends were found in the other DSS-treated groups of mice, but these changes in microbiota composition did not reach statistical significance. An increase in the iron content of the diet changed the microbiota after colitis was induced with DSS, which was not observed in the standard or lower dietary iron groups. A shifting balance within the intestinal microbiota could alter host immune response and open niches for the establishment of key environmental-shaping bacteria in the intestine, for example, the significant decrease in numbers of beneficial Firmicutes could create an opportunity for, and encourage the growth of potential gut pathogens ^45^. Bacteria species within the Firmicutes phylum are predominant in the generation of short-chain fatty acids, particularly butyrate, from dietary metabolism of insoluble fibre, resistant starches and fermentable soluble fibres (non-starch polysaccharides), ^46^, thereby providing a key anti-inflammatory effectors to ameliorate animal models of colitis

This is the first study to use models of colitis to contemporaneously assess the influence of dietary iron content on both disease activity and the microbiome. It emphasises the detrimental effects of both halving and doubling the amount of iron in the diet on a murine model of IBD. The diet with double the standard level of iron (400 ppm) led to key changes in the microbiome and this would imply that these changes observed were not simply driven by the severity of inflammation, but rather that lumenal free iron can also contribute to the complex interaction of factors that lead to the development of a dysbiotic state as has frequently been observed in IBD. There is more to understand how all sources of luminal iron influence IBD. Furthermore, work is needed to outline the physiological impact on the gut microbiota resultant from increased availability of luminal iron and how this may affect bacterial phyla and diversity. Future intervention studies in humans will be invaluable to further define the complex effects of different doses of therapeutic oral iron on the human gut microbiota, particularly to understand the metabolic consequences of observed phyla changes.

“Authors’ contributions.”
**Awad Mahalhal** [conceived and designed research, performed experiments, analysed data, interpreted results of experiments, prepared figures, drafted manuscript, edited and revised manuscript and approved the final version of manuscript]
**Jonathan M. Williams** [interpreted results of experiments, drafted manuscript, approved the final version of manuscript]
**Sophie Reade** [conceived and designed research]
**Nicholas Ellaby** [analysed data]
**Carrie A. Duckworth** [conceived and designed research, edited and revised manuscript, approved the final version of the manuscript]
**Michael D. Burkitt** [analysed data, edited and revised manuscript, approved the final version of manuscript]
**Xuan Liu** [analysed data]
**Georgina L. Hold** [analysed data, prepared figures, drafted manuscript, edited and revised manuscript, approved the final version of manuscript]
**Barry J. Campbell** [conceived and designed research, interpreted results of experiments, edited and revised manuscript, approved the final version of manuscript]
**D. Mark Pritchard** [conceived and designed research, edited and revised manuscript, approved the final version of the manuscript
**Chris S. Probert** [conceived and designed research, analysed data, interpreted results of experiments, edited and revised manuscript, approved the final version of manuscript].

